# K_v_1.3 induced hyperpolarisation and Ca_v_3.2-mediated calcium entry are required for efficient Kaposi’s sarcoma-associated herpesvirus lytic replication

**DOI:** 10.1101/2021.09.10.459757

**Authors:** Holli Carden, Mark L. Dallas, David J. Hughes, Jonathan D. Lippiat, Jamel Mankouri, Adrian Whitehouse

## Abstract

Understanding the host factors critical for Kaposi’s sarcoma-associated herpesvirus (KSHV) lytic replication can identify new targets for therapeutic intervention. Using pharmacological and genetic silencing approaches, we reveal for the first time that KSHV requires a B cell expressed voltage-gated K^+^ channel, K_v_1.3, to enhance lytic replication. We show that the KSHV replication and transcription activator (RTA) protein upregulates K_v_1.3 expression, leading to enhanced K^+^ channel activity and hyperpolarisation of the B cell membrane. Enhanced K_v_1.3 activity then promotes intracellular Ca^2+^ influx through Ca_v_3.2, a T-type Ca^2+^ channel, leading to the Ca^2+^ driven nuclear localisation of NFAT and the subsequent NFAT1-responsive gene expression. Importantly, KSHV lytic replication and infectious virion production could be inhibited by both K_v_1.3 and Ca_v_3.2 blockers or through K_v_1.3 silencing. These findings provide new mechanistic insight into the essential role of host ion channels during KSHV infection and highlight K_v_1.3 and Ca_v_3.2 as new druggable host factors that are key to the successful completion of KSHV lytic replication.

## Introduction

Kaposi’s sarcoma-associated herpesvirus (KSHV) is a gamma 2-herpesvirus directly linked to the development of Kaposi’s sarcoma (KS), a highly vascular tumour of endothelial lymphatic origin, and several other AIDs-associated malignancies including primary effusion lymphoma (PEL) and some forms of multicentric Castleman’s disease (MCD)^1–4^. KSHV exhibits a biphasic life cycle consisting of latent persistence or lytic replication. In contrast to other oncogenic herpesviruses in which latent gene expression drives tumorigenesis, both the latent and lytic replication phases are essential for KSHV-mediated tumorigenicity^5^. Latency is established in B cells and in the tumour setting, where viral gene expression is limited to the latency-associated nuclear antigen (LANA), viral FLICE inhibitory protein, viral cyclin, kaposins and several virally-encoded miRNAs^6–8^. Upon reactivation, KSHV enters the lytic replication phase, leading to the highly orchestrated expression of more than 80 viral proteins that are sufficient for the production of infectious virions^9,10^. In KS lesions, most infected cells harbour the virus in a latent state. However, a small proportion of cells undergo lytic replication that leads to the secretion of angiogenic, inflammatory and proliferative factors that act in a paracrine manner on latently-infected cells to enhance tumorigenesis^11^. Lytic replication also enhances genomic instability^12^ and sustains KSHV episomes in latently-infected cells that would otherwise be lost during cell division^13^. The ability to inhibit the lytic replication phase therefore represents a therapeutic intervention strategy for the treatment of KSHV-associated diseases^14,15^.

The transition from latent infection to lytic replication is controlled by both host and viral factors^16,17^. These factors converge on the regulation of the latency associated nuclear antigen (LANA) and the master regulator of the latent-lytic switch, KSHV replication and transcription activator (RTA) protein^18^. Notably, agents that mobilize intracellular Ca^2+^ can induce the expression of KSHV-RTA and enhance KSHV reactivation and lytic replication^19^, however this activity can be blocked with inhibitors of calcineurin-dependent signal transduction^20^. Cytoplasmic concentrations of Ca^2+^ are regulated by a network of ion channels and transporters^21^. Ion channels are multi-subunit, pore-forming membrane proteins that control the rapid and selective passage of ions across the plasma membrane and the membranes of subcellular organelles^22^. As such, ion channels have a wide variety of roles in controlling the ion homeostasis of the cell and its organelles, action potential firing, membrane potential and cell volume. Given this wide range of functions and their ubiquitous nature, dysregulation of ion channels have been implicated in a variety of disorders and diseases known as channelopathies^23^ and may also play an important role in enhancing cell proliferation and invasion of tumour cells.

Several stages of virus replication cycles, including virion entry, virus egress and the maintenance of an environment conducive to virus replication are in-part, dependent on the ability of virus proteins to manipulate ion channel activity^23,24^. Most notably, the pharmacological modulation of virus-targeted ion channels has been shown to impede virus replication, highlighting ion channels as promising candidates for host targeted anti-viral therapeutics. A catalogue of ion channel-blocking drugs have also been shown to possess antiviral activity, some of which are in widespread human use for ion channel-related diseases, highlighting new potential for drug repurposing. Amongst herpesviruses, varicella-zoster virus and herpes simplex virus have been shown to activate voltage-gated Na^+^ channels and voltage-gated Ca^2+^ channels^25,26^. However, a specific role for host cell ion channels during the lytic replication stage of KSHV or any herpesvirus have yet to be fully defined. B lymphocytes, the primary site of KSHV latent infection, are regulated by a network of transporters and ion channels that control the cytoplasmic concentrations of calcium (Ca^2+^), magnesium (Mg^2+^) and zinc (Zn^2+^), which act as important second messengers to regulate critical B cell effector functions^27^. The repertoire of ion channels in B cells include potassium (K^+^) channels, Ca^2+^ channels, P2X receptors and transient receptor potential (TRP) channels, in addition to Mg^2+^ and Zn^2+^ transporters. To-date, a role for these channels during KSHV infection has not been described.

Here, we performed a systematic analysis of the role of host ion channels during the KSHV lytic replication phase in human B cells to reveal new avenues for host-directed therapeutic intervention. Using a combination of electrophysiological and biochemical approaches, we show that KSHV activates a voltage-gated K^+^ channel K_v_1.3, the pharmacological and genetic silencing of which inhibits KSHV lytic replication. We further define the mechanism for this dependence by showing that K_v_1.3 activation leads to hyperpolarisation induced Ca^2+^ influx through a T-type Ca^2+^ channel Ca_v_3.2, which enhances the nuclear localisation of NFAT1, which in turn is required to drive virus replication. We therefore reveal for the first time the essential role of K_v_1.3 and Ca_v_3.2 channels in the lytic replication cycle of a herpesvirus.

## Material and Methods

### Cell culture, antibodies and reagents

TREx-BCBL-1-RTA cells (kindly provided by Prof. Jae Jung, University of Southern California) are a BCBL-1-based primary effusion lymphoma (PEL) B cell line engineered to express exogenous Myc-tagged RTA upon addition of doxycycline, triggering reactivation of the KSHV lytic cycle. A549 and HEK-293T cell lines were purchased from the American Type Culture Collection (ATCC). U87 cells (kindly provided by Prof. J. Ladbury, University of Leeds) are a human brain glioblastoma astrocytoma cell line. A549, U87 and HEK-293T cells were grown in DMEM (Life Technologies) supplemented with 10% foetal calf serum (FCS) (Life Technologies) and 1% penicillin/streptomycin (P/S). TREx BCBL1-RTA cells were grown in RPMI 1640 medium (Life Technologies) supplemented with 10% FCS and 1% P/S, and maintained under hygromycin B (Life Technologies) selection (100 μg/ml). Reactivation into the lytic cycle was induced using 2 μg/ml doxycycline hyclate (Sigma) as previously described^28^. All cells were maintained at 37°C in a humidified incubator with 5% CO_2_. pEGFP-N1 was obtained from Clontech, pRTA-EGFP and pORF57-GFP have been described previously^29–31^. Rabbit antibodies anti-Sp1 (EPR22648-50), anti-Lamin B1 and anti-NFAT1 were purchased from Abcam. Mouse monoclonal antibodies to GAPDH (1E6D9) and KSHV ORF57 (207.6) were obtained from Proteintech and Santa Cruz, respectively. Mouse monoclonal anti-c-Myc (9E10) and rabbit polyclonal anti-K_v_1.3 antibodies were obtained from Sigma. KSHV ORF59 and KSHVGFP ORF65 were kind gifts from Prof. Britt Glaunsinger (University of California, Berkeley) and Prof. Shou-Jiang Gao (University of Pittsburgh), respectively.

(2R/S)-6-prenylnaringenin, Blood Depressing Substance I and ShK-Dap^22^ were obtained from Bio-Techne. Mibefradil and pimozide were purchased from Cambridge Bioscience. 4-aminopyridine, calcium ionophore A23187, charybdotoxin, margatoxin, nifedipine, quinine hydrochloride dihydrate, SKF-96365, tetraethylammonium chloride and TRAM-34 were obtained from Sigma-Aldrich. Flunarizine, Cyclosporin A, Mithramycin A and KCl were purchased from Alfa Aesar, Fluorochem Limited, Insight Biotechnology and ThermoFisher, respectively. Human foetal brain (HFB) cDNA library was purchased from Invitrogen. Oligonucleotide primer sequences are available upon request. All primers were purchased from Sigma (UK).

### Lentivirus-based shRNA Knockdown

Lentiviruses were generated by transfection of HEK-293T cells seeded in 12-well plates using a three-plasmid system, as previously described^28^. Per 6-well, 4 μl of lipofectamine 2000 (Thermo Scientific) were used together with 1 μg of pLKO.1 plasmid expressing shRNA against the protein of interest (Dharmacon), 0.65 μg of pVSV.G, and 0.65 μg psPAX2. pVSV.G and psPAX2 were a gift from Dr. Edwin Chen (University of Westminster, London). Eight hours post-transfection, media was changed with 2 mL of DMEM supplemented with 10% (v/v) FCS. 500,000 TREx BCBL1-Rta cells in 6 well plates were infected by spin inoculation with the filtered viral supernatant for 60 min at 800 × g at room temperature, in the presence of 8 μg/mL of polybrene (Merck Millipore). Virus supernatants were removed 7 h post-spin inoculation and cells were maintained in fresh growth medium for 48 h prior to selection in 3 μg/mL puromycin (Sigma-Aldrich). Stable cell lines were generated after 8 days of selection.

### Transient Transfections

Plasmid transfections were performed using Lipofectamine 2000 (Life Technologies), at a ratio of 2 ug plasmid to 1 ul Lipofectamine in 100 ul opti-MEM. Transfection media was incubated at room temperature for 15 minutes before 1× 10^6^ cells were treated, dropwise. Cells were harvested after 24 hours.

### Immunofluorescence

Cells were cultured overnight on poly-L-lysine (Life Technologies) coated glass coverslips in 24-well plates. Cells were fixed with 4% paraformaldehyde (Calbiochem) for 10 min and permeabilised with 0.1% Triton X-100 for 20 min as previously described^32^. Cells were blocked in PBS containing 1% BSA for 1 h at 37°C and labelled with primary antibodies for 1 h at 37°C. Cells were washed five times with PBS and labelled with appropriate secondary antibodies for 1 h at 37°C. Cells were washed five times with PBS and mounted in VECTASHIELD containing DAPI (Vector Labs). Images were obtained using a Zeiss LSM700 Inverted Microscope confocal microscope and processed using ZEN 2009 imaging software (Carl Zeiss) as previously described^29^.

### Electrophysiology

TREx BCBL1-RTA cells seeded onto poly-L-lysine (Life Technologies) coated glass coverslips and were transferred to a recording chamber, containing 140 mM NaCl, 5 mM KCl, 2 mM MgCl_2_, 10 mM HEPES-NaOH, pH 7.2, 2 mM CaCl_2_, 10 mM glucose, and mounted on the stage of a Nikon Eclipse inverted microscope. Patch pipettes (5–8 MΩ) were filled with a solution consisting of: 140 mM KCl, 5 mM EGTA, 2 mM MgCl_2_, 1 mM CaCl_2_, 10 mM HEPES KOH, pH 7.2, 10 mM glucose. Voltage-clamp recordings were performed using a HEKA EPC-10 integrated patch clamp amplifier controlled by Patchmaster software (HEKA). Series resistance was monitored after breaking into the whole cell configuration. To examine K^+^ currents, a series of depolarizing steps were performed from −100 to +60 mV in 10 mV increments for 100 ms. Resting membrane potential was measured using the current clamp mode of the amplifier. Results are shown as the mean ± SEM of n number of individual cells. Statistical analysis was performed using an unpaired Student’s T test. p<0.05 was considered statistically significant.

### Flow Cytometry

Bis-(1,3-Dibutylbarbituric Acid) Trimethine Oxonol (DiBAC_4_(3)) and Fura Red (both ThermoFisher) were added to cells at a final concentration of 1 μM in RPMI-media. Cells were incubated at 37°C with Fura Red for 30 min or DiBAC_4_(3) for 5 min and washed in PBS. Cells were analysed on a CytoFLEX Flow Cytometer (Beckman). Data were quantified using FlowJo software as previously described^33^ (Tree Star, Ashland, OR, USA).

### Proliferation (MTS) assays

Cellular viability was determined using non-radioactive CellTiter 96 AQueous One Solution Cell Proliferation Assay (MTS) reagent (Promega), according to the manufacturer’s recommendations^29^. Briefly, TREx BCBL1-RTA cells (^∼^20,000) were seeded in triplicate in a flat 96-well tissue culture plates (Corning) and treated with the indicated inhibitors for 24 h. CellTiter 96 AQueous One Solution Reagent was added to the cells for 1 h at 5% CO_2_, 37°C. Absorbances were measured at 490 nm using an Infinite plate reader (Tecan).

### Two-step quantitative reverse transcription PCR (qRT-PCR)

Total RNA was extracted using the Monarch^®^ Total RNA Miniprep Kit (New England Biolabs) as per the manufacturer’s protocol. RNA (1 μg) was diluted in a total volume of 16 μl nuclease-free water, and 4 μl LunaScript RT SuperMix (5X) (New England Biolabs) was added to each sample. Reverse transcription was performed using the protocol provided by the manufacturer. cDNA was stored at - 20°C, RNA was stored at −80°C. Quantitative PCR (qPCR) reactions (20 μl) included 1X SensiMix SYBR green master mix (Bioline), 0.5 μM of each primer and 5 μl template cDNA (used at 1:200 dilution in RNase-free water). Cycling was performed in a RotorGene Q instrument (Qiagen), using a previously described cycling programme^29^. Relative gene expression was calculated using the ΔΔCT method, as previously described^29^. Statistical significance was validated using a Student’s t-test.

### Chromatin immunoprecipitation (ChIP)

Formaldehyde-crosslinked chromatin was prepared using the Pierce Chromatin Prep Module (Thermo Scientific) following the manufacturer’s protocol. Cells (2 × 10^6^) were digested with six units of micrococcal nuclease (MNase) per 100 μl of MNase Digestion buffer in a 37°C water bath for 15 min. These conditions resulted in optimal sheared chromatin with most fragments ranging from 150 –300 base pairs in size. Immunoprecipitations were performed using EZ-ChIP kit (Millipore) kits overnight at 4°C and contained 50 μl of digested chromatin (2 × 10^6^ cells), 450 μl of ChIP dilution buffer and 1.5 μg of RNAPII antibody (clone CTD4H8) (Millipore) or isotype antibody, normal mouse IgG (Millipore). qPCR reactions were performed using either 2 μl of immunoprecipitated DNA or 2 μl of input DNA as template.

### Immunoblotting

Protein samples were separated on SDS-PAGE gels as previously described^34^, and transferred to nitrocellulose membranes (Amersham) via semi-dry transfer using a Trans-Blot® Turbo™ blotter (BioRad). Membranes were blocked in TBS + 0.1% Tween 20 and 5% dried skimmed milk powder and probed with relevant primary antibodies followed by horseradish peroxidase (HRP)-conjugated polyclonal goat anti-mouse and polyclonal goat anti-rabbit secondary antibodies (Dako). Membranes were treated with EZ-ECL (Geneflow) and imaged using a G-Box (Syngene).

## Results

### K^+^ channels are required for efficient KSHV reactivation

K^+^ channels represent the largest family of ion channels with over 70 genes identified in the human genome^35^. We first sought to determine if their activity is required for efficient KSHV lytic replication. Here virus reactivation assays were performed in the presence of potassium chloride (KCl) to collapse cellular K^+^ channel gradients, or the broad spectrum K^+^ channel blockers, barium chloride (BaCl_2_), tetraethylammonium (TEA) and quinidine (Qn). All were assessed at non-toxic concentrations (Supp Fig 1). KSHV reactivation was assessed in TREx BCBL1-RTA cells, a latently infected KSHV B-lymphocyte cell line that express a Myc-tagged viral RTA under the control of a doxycycline-inducible promoter. Upon analysis, TREx BCBL1-RTA cells reactivated for 24 h in the presence of each K^+^ channel inhibitor showed a drastic reduction in the expression early ORF57, delayed early ORF59 and the late minor capsid (mCapsid) ORF65 proteins (**Fig 1a**). No such reduction was observed in the expression of Myc-RTA or GAPDH, highlighting specific effects on lytic replication as opposed to dox-induced RTA induction. These data indicate a requirement for K^+^ channel function during the KSHV lytic replicative cycle.

**Figure 1.**
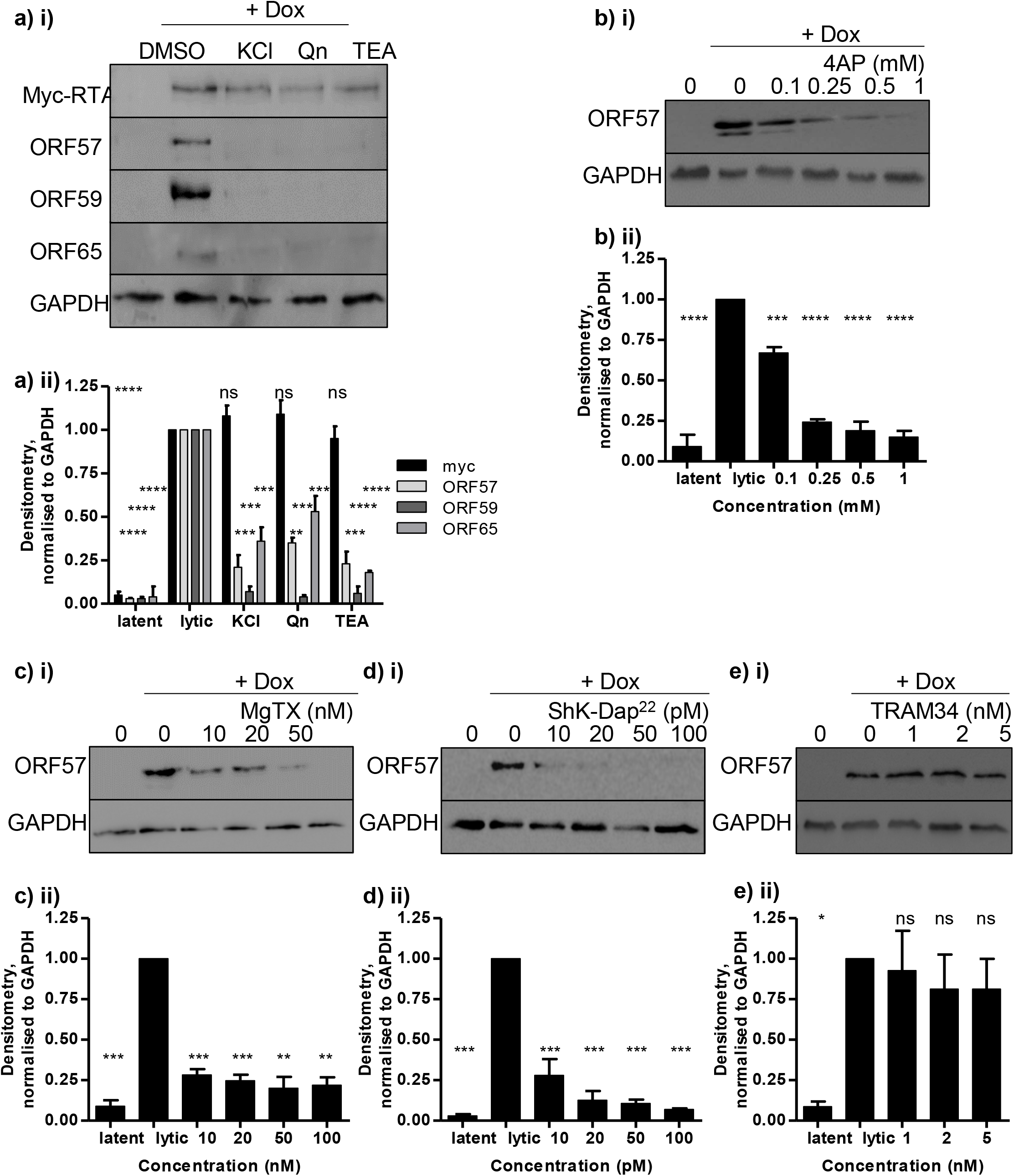
K+ channels are required for efficient KSHV lytic replication. TREx BCBL1-RTA cells remained unreactivated or were pre-treated with non-cytotoxic concentrations of (A) general K^+^ inhibitors, 25 mM KCl, 100 μM Qn and 10 mM TEA, or increasing amounts of (B) 4AP (C) MgTX, (D) ShK-Dap^22^ and (E) TRAM34 for 45 minutes prior to reactivation with doxycycline hyclate for 48 hr. (i) Cell lysates were then probed with ORF57-, ORF59- or ORF65-specific antibodies. GAPDH was used as a measure of equal loading (ii) Densitometry quantification of immunoblots was carried out using the Image J software and is shown as a percentage relative to the loading control, GAPDH. Data analysed using three replicates per experiment, n = 3 and statistical analysis using a two-tailed t-test with unequal variance, ** = p<0.01, **** = p<0.001, **** = p<0.0001.

We next investigated the molecular identity of the specific K^+^ channel(s) required for KSHV lytic replication to reveal more specific drug targets. K^+^ channels can be divided into subfamilies of voltage-gated K^+^ channels (K_v_), calcium-activated K^+^ channels (K_Ca_), inwardly rectifying K^+^ channels (K_ir_) and two-pore domain K+ channels (K2P) channels. We found that treatment with 4-aminopyridine (4-AP), a non-selective K_v_ blocker, led to a concentration-dependent reduction in lytic replication (**Fig 1b**), suggestive of a role for K_v_ channels during lytic induction. Electrophysiological studies have identified an array of functional K_v_ channels expressed within B lymphocytes, with a member of the *Shaker* related family, K_v_1.3, most extensively characterised^36^. When specific K_v_1.3 blockers margatoxin (MgTX) and ShK-Dap^22^ were included in our reactivation assays, a concentration-dependent reduction of lytic gene expression was observed, implicating a role for this channel during lytic KSHV replication (**Fig 1c-d**). In contrast, TRAM-34, a blocker of B lymphocyte K_Ca_3.1 channels, showed no effect on lytic KSHV protein production (**Fig 1e**).

To confirm a role for K_v_1.3 during KSHV lytic replication, TREx BCBL1-RTA cells were transduced with lentivirus-based shRNAs to effectively deplete K_v_1.3 by over 85% (**Fig 2a-b**). Reactivation assays showed that K_v_1.3 silencing led to a significant reduction in ORF57, ORF59 and ORF65 proteins compared to scrambled controls (**Fig 2c**). To examine whether the depletion of K_v_1.3 also influenced infectious virus production, supernatants of reactivated TREx BCBL1-RTA cells were used to re-infect naive cells and KSHV infection was determined by qRT-PCR. Cells reinfected with supernatants from K_v_1.3 depleted cells showed a ^∼^85% reduction in viral mRNA compared to control cells (**Fig. 2d**). Together, these data confirmed that KSHV requires the function of B cell K_v_1.3 channels to undergo efficient lytic replication and infectious virus production.

**Figure 2.**
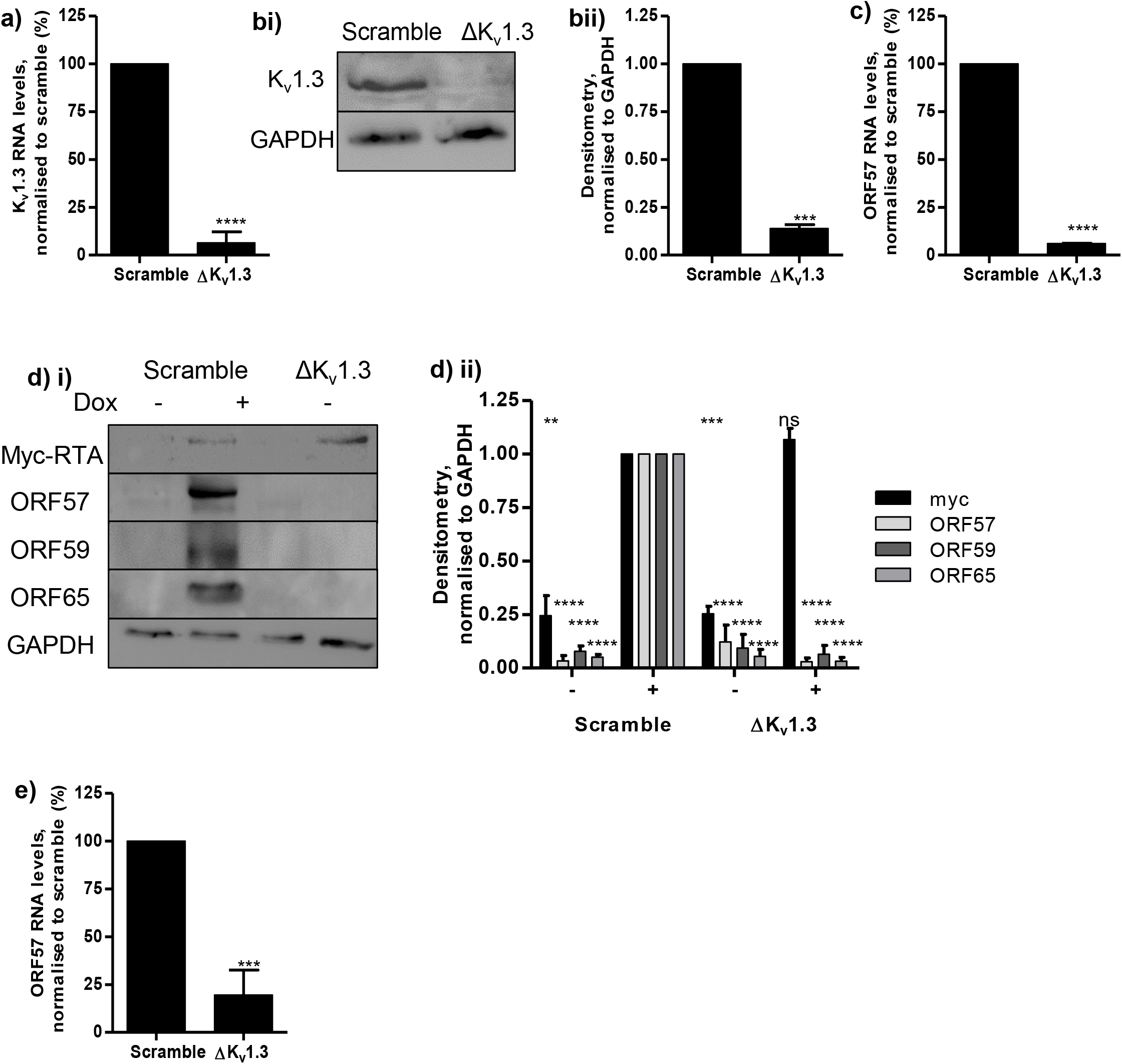
K_v_1.3 is required for efficient KSHV lytic replication. TREx BCBL1-RTA cells were stably transduced with lentivirus expressing scrambled or K_v_1.3-specfic shRNAs. Scramble and K_v_1.3-depleted cells lines were reactivated with doxycycline hyclate for 24 hr. (A) Total RNA was extracted and relative transcript levels were analysed by qRT-PCR using GAPDH as a reference. Fold change was determined by ΔΔCt and statistical significance analysed using a non-paired t-test, **** = p<0.0001. B and C (i) Cell lysates were probed with (B) K_v_1.3- or (C) ORF57-, ORF59- or ORF65-specific antibodies and GAPDH used as a measure of equal loading (ii) Densitometry quantification of immunoblots was carried out using the Image J software and is shown as a percentage relative to the loading control, GAPDH. Data analysed using three replicates per experiment, n = 3 and statistical analysis using a two-tailed t-test with unequal variance, *** = p<0.001, **** = p<0.0001. (D) Scramble or K_v_1.3-depletetd TREx BCBL1-RTA cells were reactivated for 72 h. The culture medium was then incubated for 24 h with HEK-293T cells followed by total RNA extraction and qRT-PCR. Results show the mean of three biological replicates with error bar as standard deviation, *** = p<0.001.

### KSHV enhances K_v_1.3 expression and activity

As KSHV is dependent on K_v_1.3 to complete its lytic replication cycle, its ability to modulate K_v_1.3 activity was next investigated. qRT-PCR and immunoblotting analysis showed that K_v_1.3 expression increased in TREx BCBL1-RTA cells undergoing lytic replication compared to latent cells (**Fig. 3a**). To elucidate whether the increase in K_v_1.3 expression led to enhanced K^+^ efflux mediated by K_v_1.3 channels during lytic replication, whole-cell patch clamp analysis was performed. Electrophysiological recordings revealed a voltage-gated outward K^+^ current present in latent TREx BCBL1-RTA cells that was significantly enhanced in cells undergoing lytic replication (**Fig. 3b**). To conclusively determine that K_v_1.3 channels were responsible for these changes, recordings were repeated in the presence of ShK-Dap^22^, which led to a dramatic inhibition of the K^+^ current (**Fig. 3b**). A similar reduction was observed in cells depleted for K_v_1.3 using lentivirus-based shRNAs, compared to scrambled controls (**Fig 3c**). Notably, we also observed that reactivated TREx BCBL1-RTA cells exhibited a significantly more hyperpolarised membrane compared to latent cells (**Fig. 3d**), which was reversed upon K_v_1.3 depletion (**Fig. 3e**). Membrane hyperpolarisation was also confirmed using a membrane potential-sensitive dye, bis (1,3-dibutylbarbituric acid) trimethine oxonol; DiBAC_4_(3). Results showed a time-dependent decrease in fluorescence intensity in lytic cells, consistent with enhanced membrane hyperpolarization (**Fig 3e**). In contrast, addition of the calcium ionophore A23187, which induces depolarisation, enhanced DiBAC4(3) fluorescence. Moreover, no reduction in DiBAC4(3) fluorescence was observed in K_v_1.3 depleted cells (**Fig 3f**). Together these results demonstrate that KSHV lytic replication increases K_v_1.3 expression, resulting in an enhanced K_v_1.3 currents and hyperpolarisation during lytic KSHV replication.

**Figure. 3.**
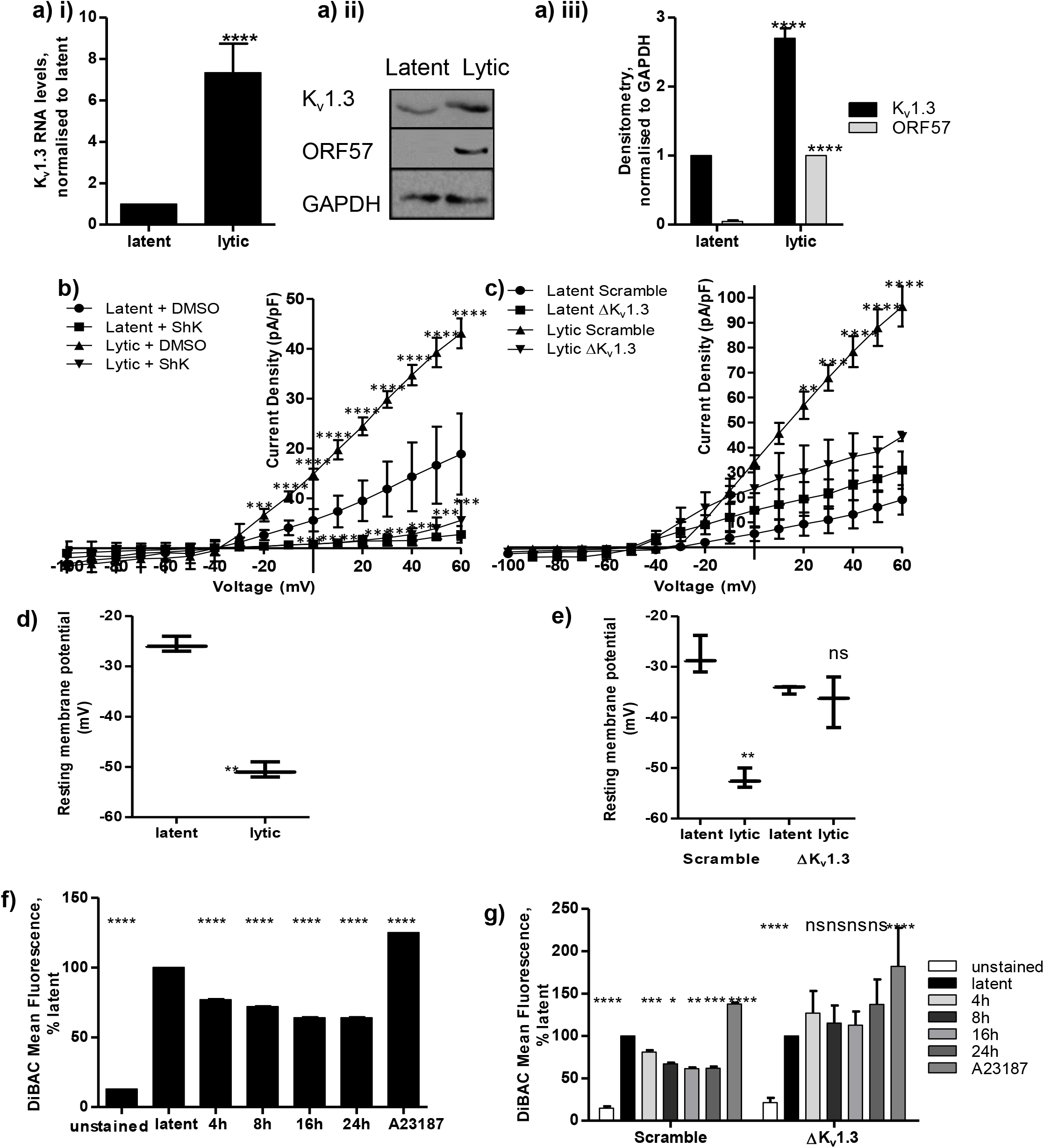
Increased K_v_1.3 currents during lytic KSHV replication. (A) TREx BCBL1-RTA cells remained unreactivated or were reactivated with doxycycline hyclate for 24 hr. (i) Total RNA was extracted and relative K_v_1.3 transcript levels were analysed by qRT-PCR using GAPDH as a reference. Fold change was determined by ΔΔCt and statistical significance analysed using a non-paired t-test, **** = p<0.0001. (ii) Cell lysates were probed with K_v_1.3-specific antibodies and GAPDH used as a measure of equal loading (iii) Densitometry quantification of immunoblots was carried out using the Image J software and is shown as a percentage relative to the loading control, GAPDH. Data analysed using three replicates per experiment, n = 3 and statistical analysis using a two-tailed t-test with unequal variance, *** = p<0.001, **** = p<0.0001. Mean current density voltage relationships for K^+^ currents (n=5 for all populations) from (B) unreactivated and reactivated TREx BCBL1-RTA at 16 hr; cells were pre-treated for 24 hours with DMSO control or 100 pM ShK-Dap^22^ and (C) Scramble and K_v_1.3-depleted cells lines which remained unreactivated or were reactivated with doxycycline hyclate for 24 hr. Pooled data highlighting resting membrane potentials in (D) latent and lytic TREx BCBL1-RTA cells or (E) Scramble and K_v_1.3-depleted cells lines. Membrane polarisation of TREx BCBL1-RTA cells was measured by Flow cytometry after a 5 min incubation with DiBAC4(3) (F) Unreactivated and reactivated TREx BCBL1-RTA at 24 hr or (G) Scramble and K_v_1.3-depleted cells lines.

### KSHV RTA mediates the upregulation of K_v_1.3 during lytic replication

We next investigated the mechanism by which KSHV enhances K_v_1.3 currents. Given that membrane hyperpolarisation was observed as early as 4 h post-reactivation, we examined whether the KSHV early proteins were sufficient to induce K_v_1.3 expression and activation. A549 and U87 cells were transiently transfected with control GFP, RTA-GFP or ORF57-GFP expression constructs and K_v_1.3 transcript levels were assessed by qRT-PCR at 24 h post-transfection. We found that RTA-GFP alone was sufficient to induce K_v_1.3 expression at the transcript level (**Fig 4a-b**), confirming KSHV RTA transcriptional activator as the direct inducer of K_v_1.3 expression.

**Figure 4.**
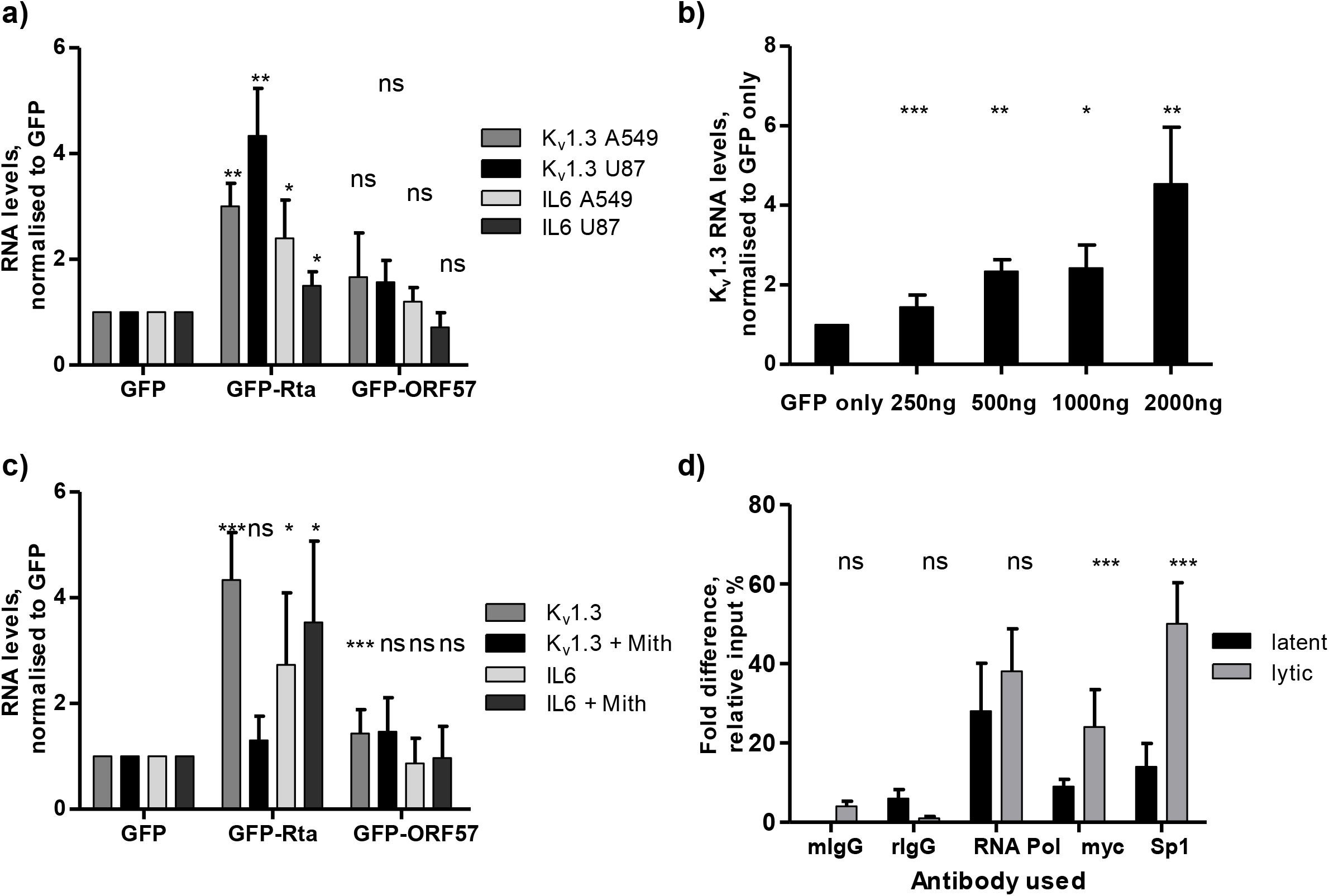
KSHV RTA upregulates K_v_1.3 expression. A549 and U87 cells were transfected with either (A) GFP, RTA-GFP or ORF57–GFP expression construct for 24 hr, Dose-dependent increase of RTA-GFP for 24 hr or (C) GFP, RTA-GFP or ORF57–GFP in the presence of the Sp1 inhibitor, Mithramycin A. Total RNA was extracted and relative K_v_1.3 or IL-6 transcript levels were analysed by qRT-PCR using GAPDH as a reference. Fold change was determined by ΔΔCt and statistical significance analysed using a non-paired t-test, * = p<0.05, ** = p<0.01, *** = p<0.001. (D) ChIP assays were carried out on unreactivated versus reactivated TREx BCBL1-RTA cells using antibodies specific to SP-1, RNAPII, myc-RTA or mouse and rabbit IgG control antibodies. PCR amplification was carried out on the immunoprecipitates using primers against the K_v_1.3 promoter region to determine association. The average of three independent experiments is shown with error bars as standard deviation, *** = p<0.001.

Specificity Protein (Sp) 1 functions as a co-adapter for RTA-mediated transactivation and has been shown to regulate K_v_1.3 expression^37^. To further dissect the relationship between KSHV-RTA and K_v_1.3, we examined a potential cooperative role for Sp1 during the upregulation of K_v_1.3 during lytic replication. RTA-GFP transfections were performed in the presence of Mithramycin A, a selective Sp1 inhibitor that displaces Sp1 binding from its target promoter^38^. Results showed Mithramycin A treatment supressed the RTA-mediated increase in K_v_1.3 expression (**Fig 4c**), but had little effect on the RTA-mediated upregulation of the IL-6 promoter, suggestive of an in-direct mechanism whereby Sp1 recruits RTA to the K_v_1.3 promoter. ChIP assays further confirmed an association of both RTA and Sp1 with the K_v_1.3 promoter, which significantly increased during lytic replication (**Fig 4d**). Together, these data reveal KSHV RTA as the key driver of K_v_1.3 expression during the KSHV lytic cycle.

### K_v_1. 3 induced membrane hyperpolarisation provides the driving force for Ca^2+^ influx required for KSHV reactivation

In B lymphocytes, K_v_1.3 maintains a hyperpolarised membrane potential that is necessary to sustain the driving force for Ca^2+^ entry. K_v_1.3 therefore indirectly modulates an array of Ca^2+^-dependent cellular processes in B cells. To assess the role of K_v_1.3 during lytic replication, we assayed Ca^2+^ influx into TREx BCBL1-RTA cells during the KSHV lytic cycle using the ratiometric Ca^2+^ dye Fura-Red and flow cytometry analysis. We observed an increase in cytoplasmic Ca^2+^ over a 24 h period of lytic reactivation (**Fig 5a**), that was absent in K_v_1.3-depleted TREx BCBL1-RTA cells compared to scrambled controls (**Fig. 5b**). Based on these data, we investigated whether Ca^2+^ influx defines the requirement of K_v_1.3 for efficient lytic replication. TREx BCBL1-RTA cells were reactivated in the absence or presence of the Ca^2+^ ionophore A23187, which led to enhanced lytic ORF57 protein levels compared to control cells (**Fig. 5c)**. We next determined whether A23187 had the ability to recover KSHV lytic replication in a K_v_1.3 depleted cell line. Notably, lytic ORF57 protein production was observed in K_v_1.3 silenced cells, suggesting that A23187 could override the dependence of KSHV on K_v_1.3 (**Fig 5d**). Together, these data suggest that Ca^2+^ influx is essential for efficient KSHV lytic replication and is induced by K_v_1.3-mediated hyperpolarisation.

**Figure 5.**
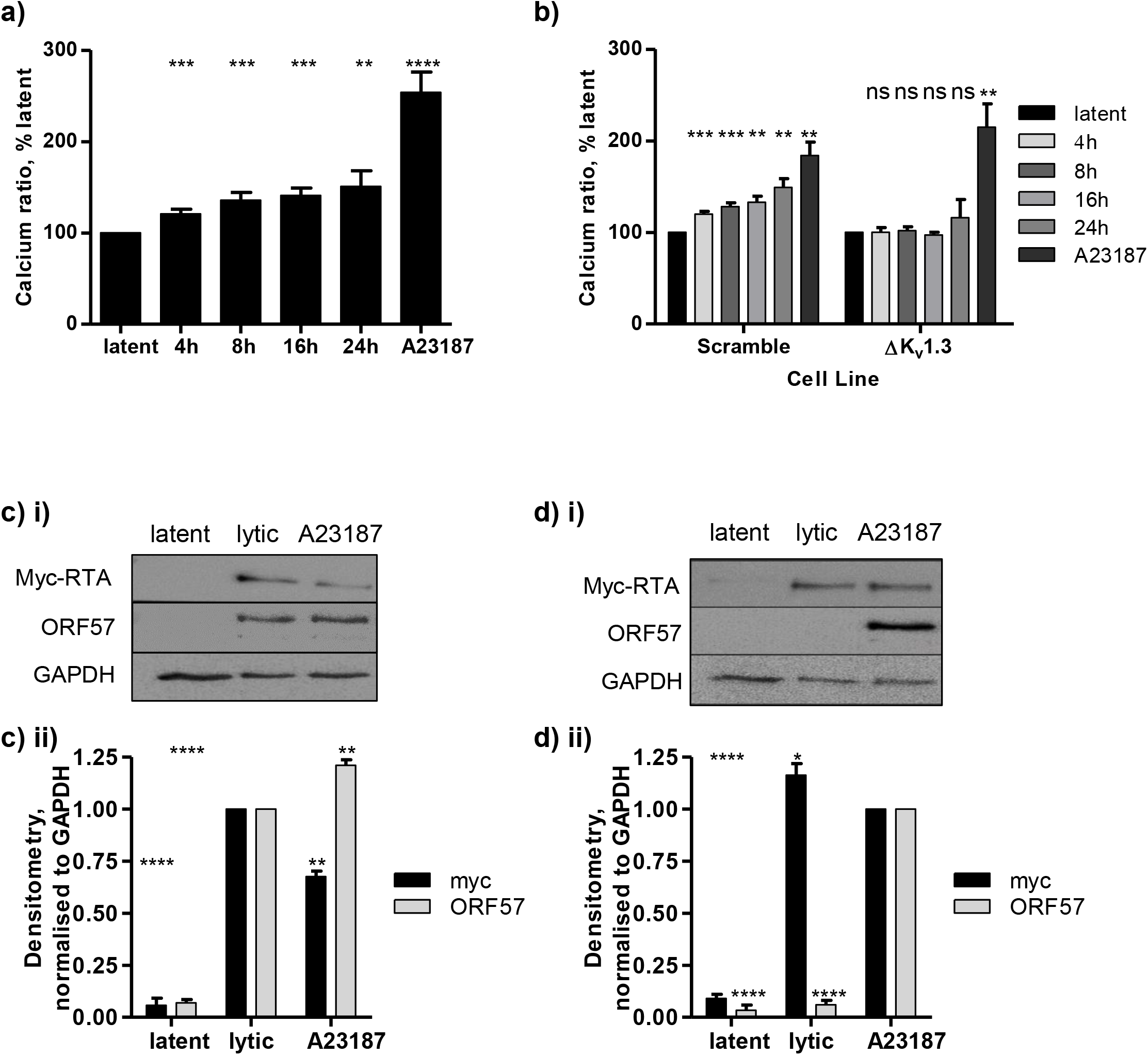
Membrane hyperpolarisation leads to Ca^2+^ influx during KSHV lytic replication. Fura Red staining of calcium ratios were measured in TREx BCBL1-RTA cells by Flow cytometry, the calcium ionophore A23187 was used as a positive control (A) Unreactivated and reactivated TREx BCBL1-RTA at 24 hr or (B) Scramble and K_v_1.3-depleted cells lines. (C) Unreactivated and reactivated TREx BCBL1-RTA at 24 hr or (D) Scramble and K_v_1.3-depleted cells lines were assessed for levels of lytic replication in the presence of the calcium ionophore A2318, (i) Cell lysates were probed with ORF57-specific antibodies and GAPDH used as a measure of equal loading (ii) Densitometry quantification of immunoblots was carried out using the Image J software and is shown as a percentage relative to the loading control, GAPDH. Data analysed using three replicates per experiment, n = 3 and statistical analysis using a two-tailed t-test with unequal variance, *** = p<0.001, **** = p<0.0001.

### KSHV-mediated Ca^2+^ influx requires the T-type voltage-gated calcium channel, Ca_v_3.2

We next sought to identify the Ca^2+^ channel(s) required for KSHV-mediated Ca^2+^ influx. Here KSHV lytic replication was assessed in presence of specific Ca^2+^ channel modulating drugs at non-cytotoxic concentations, to identify candidate Ca^2+^ channel members (**Supp Fig 1**). Results showed that SKF-96365, an inhibitor of TRP channels and STIM1, and T-type calcium channel blocking drugs dramatically reduced the levels of KSHV lytic replication (**Fig 6ai**). In contrast, Nifedipine, an L-type voltage-gated Ca^2+^ channel inhibitor had no significant effect on KSHV lytic replication (**Fig 6aii**). Additional reactivation assays performed in the presence of Mibefradil reinforced the requirement of T-type voltage-gated Ca^2+^ channels for KSHV lytic replication (**Fig 6aiii**).

**Figure 6.**
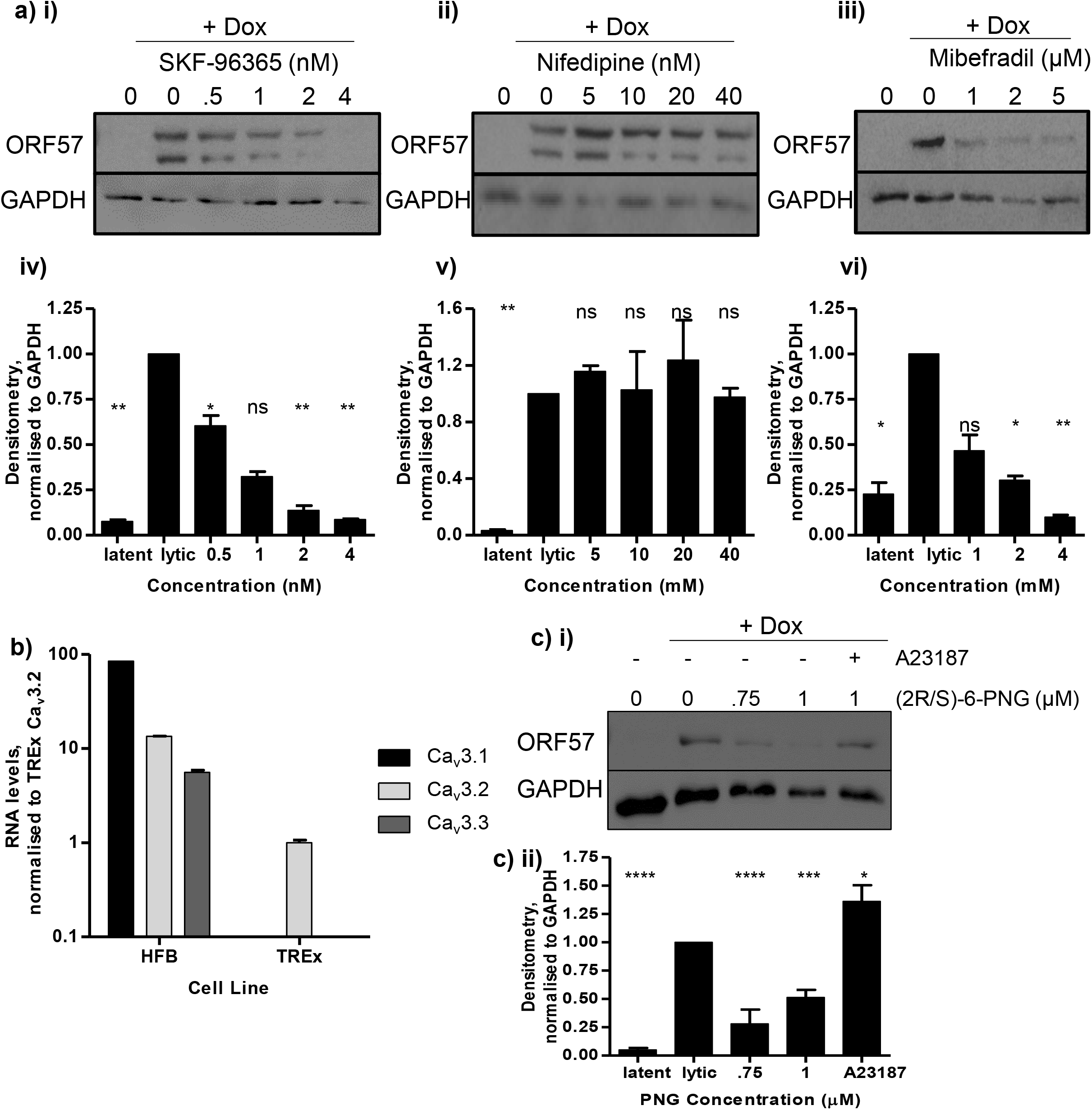
KSHV-mediated calcium influx requires the T-type voltage-gated calcium channel, Ca_v_3.2. TREx BCBL1-RTA cells remained unreactivated or were pre-treated with non-cytotoxic dose-dependent concentrations of Ca^2+^ channel inhibitors, (i) T-type and TRP channel inhibitor, SKF-96365 (ii) L-type voltage-gated inhibitor, nifedipine and (iii) T-type inhibitor, mibefradil for 45 minutes prior to reactivation with doxycycline hyclate for 48 hr. Cell lysates were then probed with ORF57-specific antibodies. GAPDH was used as a measure of equal loading. (iv-vi) Densitometry quantification of immunoblots was carried out using the Image J software and is shown as a percentage relative to the loading control, GAPDH. Data analysed using three replicates per experiment, n = 3 and statistical analysis using a two-tailed t-test with unequal variance, * = p<0.01, ** = p<0.001. T-type Ca^2+^ channel expression was analysed by qRT-PCR from total RNA was extracted from latent TREx BCBL1-RTA, and Human foetal brain cDNA using GAPDH as a reference. (C) (i) TREx BCBL1-RTA cells remained unreactivated or were pre-treated with non-cytotoxic dose-dependent concentrations of 2R/S-6-PNG for 45 minutes prior to reactivation. Cell lysates were probed with ORF57-specific antibodies. GAPDH was used as a measure of equal loading. (ii) Densitometry quantification of immunoblots was carried out using the Image J software and is shown as a percentage relative to the loading control, GAPDH. Data analysed using three replicates per experiment, n = 3 and statistical analysis using a two-tailed t-test with unequal variance, *** = p<0.001, **** = p<0.0001.

In humans, the expression of the three T-type Ca^2+^ channels, namely Ca_v_3.1, Ca_v_3.2 and Ca_v_3.3, are cell type dependent^39^. We therefore performed qRT-PCR analysis to confirm the expression profiles of each channel in TREx BCBL1-RTA cells. Human foetal brain (HFB) cDNA was included in this analysis as a positive control for all three Ca_v_3.x channel types. Analysis showed that Ca_v_3.2 is the only T-type Ca^2+^ channel expressed in TREx BCBL1-RTA cells (**Fig 6b**). To verify a role for Ca_v_3.2 during KSHV-mediated Ca^2+^ influx, KSHV lytic replication was monitored in the presence of (2R/S)-6-prenylnaringenin (PNG), a specific Ca_v_3.2 blocker^40^. The results showed a dramatic reduction of ORF57 expression, an indicator of KSHV lytic replication implicating a role for this channel during lytic KSHV replication. This was reinforced by results showing that the PNG-mediated inhibition could be rescued upon the addition of A23187 (**Fig 6c**). Together, these data reveal the requirement for Ca_v_3.2-mediated Ca^2+^ influx for efficient KSHV lytic replication in response to K_v_1.3-mediated hyperpolarisation.

### KSHV-mediated Ca^2+^ influx initiates NFAT1 nuclear localisation and NFAT1-mediated gene expression

Ca^2+^ influx can initiate multiple signalling pathways, including the serine/threonine phosphatase calcineurin and its target transcription factor NFAT (nuclear factor of activated T cells) ^21^. The phosphatase activity of calcineurin is activated through binding of the Ca^2+^-calmodulin complex, displacing the calcineurin autoinhibitory domain from the active site of the enzyme. Dephosphorylation of cytoplasmic NFAT proteins by calcineurin unmasks their nuclear localization sequences, leading to nuclear translocation and NFAT-responsive gene expression. To investigate whether KSHV-mediated hyperpolarisation and Ca^2+^ influx promote the nuclear translocation of NFAT, its nuclear/cytoplasmic distribution was compared in latent versus lytic TREx BCBL1-RTA cells using immunofluorescence analysis. Results showed that NFAT1 translocates to the nucleus in lytic cells, but remains cytoplasmic during latent infection (**Fig. 7a**). This result was confirmed by immunoblot analysis of subcellular fractions of latent and lytic TREx BCBL1-RTA cells (**Fig. 7b**). The nuclear localisation of NFAT was found to be dependent on both K_v_1.3-mediated hyperpolarisation and Ca_v_3.2-dependent Ca^2+^ influx, as it was prevented by both ShK-Dap^22^ and PNG treatment (**Fig 7a**). The nuclear localisation of NFAT1 in lytic cells was also shown to require calcineurin activity, since it was inhibited in the presence of the calcineurin inhibitor, cyclosporin A (CsA) (**Fig. 7a**). Consistent with the enhanced nuclear localisation of NFAT, we observed an increase in NFAT-responsive gene expression during KSHV lytic replication, which was reduced in the presence of ShK-Dap^22^, PNG (**Fig. 7c**), and upon K_v_1.3 depletion (**Fig. 7d**). Together, these data suggest that the KSHV-induced hyperpolarisation mediated by K_v_1.3 and the subsequent influx of Ca^2+^ through Ca_v_3.2, enhances the nuclear localisation of NFAT1 and induces NFAT-driven gene expression.

**Figure 7.**
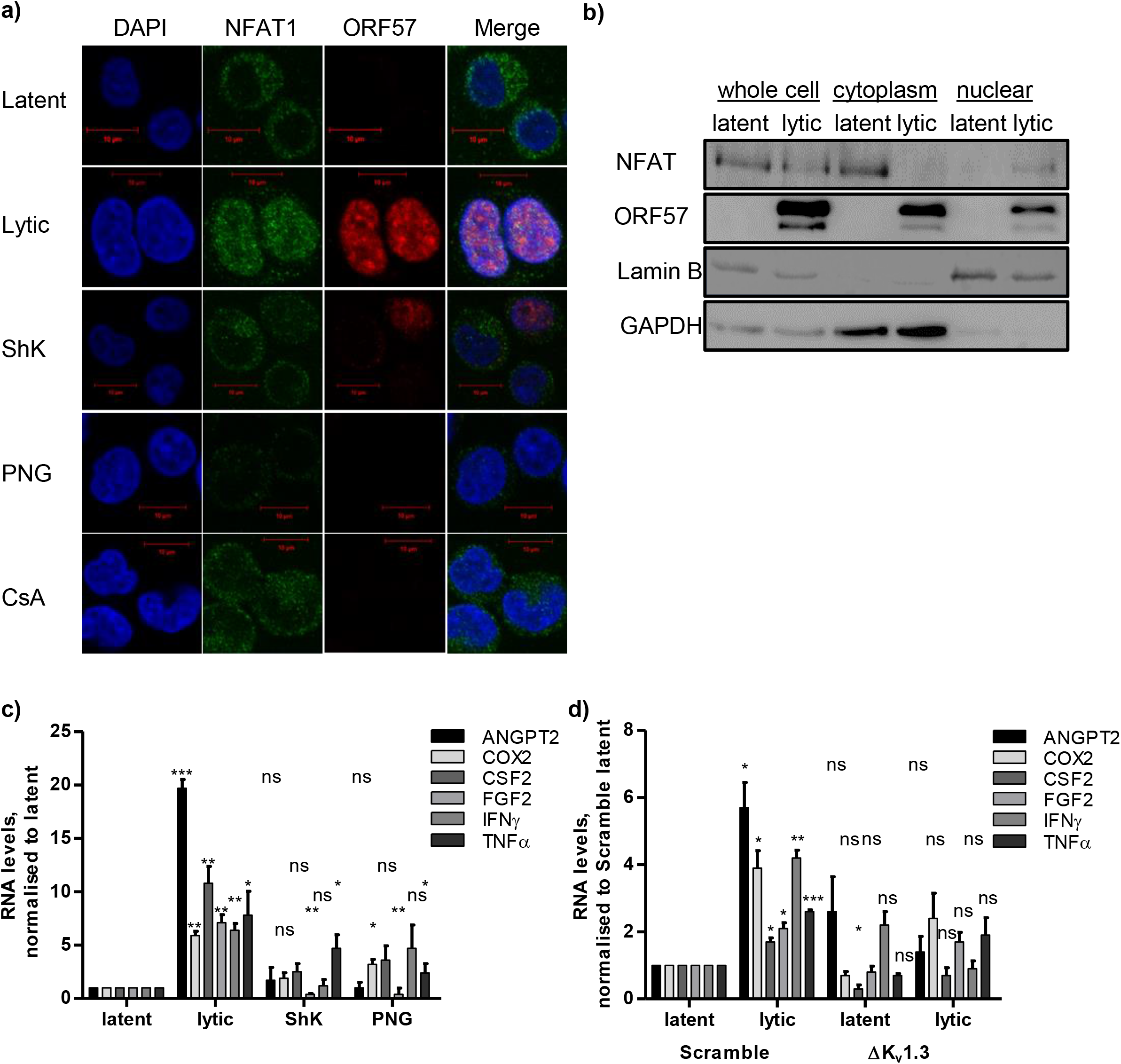
KSHV-mediated calcium influx initiates NFAT1 nuclear localisation and NFAT1-mediated gene expression. TREx BCBL1-RTA cells remained unreactivated or were pre-treated with inhibitors for 45 minutes prior to reactivation with doxycycline hyclate for 24 hr. Endogenous NFAT1 and ORF57 proteins were identified by indirect immunofluorescence using specific antibodies. (B) Cells were harvested from latent and reactivated TREx BCBL1-RTA cells at 24 hr and subcellular fractions were probed with ORF57- and NFAT-specific antibodies, GAPH and Lamin B were used as a measure of equal loading and purity of fractions. Total RNA was extracted from (C) unreactivated and reactivated TREx BCBL1-RTA cells at 24 hr, or cells pre-treated with ShK-Dap^22^ or PNG and (D) Scramble and K_v_1.3-depleted cells lines which remained unreactivated or were reactivated with doxycycline hyclate for 24 hr. Relative NFAT-responsive transcript levels were analysed by qRT-PCR using GAPDH as a reference. Fold change was determined by ΔΔCt and statistical significance analysed using a non-paired t-test, * = p<0.05, ** = p<0.01, *** = p<0.001.

## Discussion

KSHV infection is responsible for various malignancies, including KS, PEL and some cases of MCD. These diseases are highly associated with compromised immune function, and as such represent some of the most common cancers in areas of the world where HIV infection is prevalent^1^. Notably, KS is the most common cancer in many sub-Saharan countries. Therefore, understanding the molecular mechanisms that underlie KSHV biology is of the utmost importance if developing targeted therapeutic approaches. KSHV latency-associated viral proteins have been well characterised in transformation and tumourigenesis pathways; however, it is clear that KSHV also requires the lytic phase to drive tumourigenesis^14^. This is supported by a number of studies showing abrogation of KSHV gene expression impairs KSHV-associated oncogenesis. This is also emphasised by successful treatment of KS patients with drugs that inhibit KSHV replication indicate that the lytic phase is required for both the initiation of KS and the maintenance of disease^41^. Lytic genes encode angiogenic and KS growth factors which stimulate the proliferation of latently-infected cells and angiogenesis in a paracrine manner. Lytic replication can also replenish episomes lost within highly proliferating tumour cells, maintaining viral latency in select cell populations. Discovery of both the viral and cellular determinants that control lytic induction can therefore inform new therapeutic targets for anti-KSHV drug discovery. This is particularly important in light of the increasing number of AIDS-associated, Iatrogenic and classic KS cases^42^ due to the increased survival rates of AIDS patients, the higher success rates of transplant surgery, and increasing global life expectancy^43^.

Ion channels control a range of cellular processes that are known to be co-opted by viruses^23,24^. Accordingly, ion channels have emerged as druggable host targets to prevent both RNA and DNA viruses from the successful completion of their life cycles. Given the known dependence of KSHV lytic replication on Ca^2+^ signalling^20^, coupled to previous studies demonstrating the ability of VZV and HSV-1 to activate Na^+^ and Ca^2+^ family members^25,26^, we specifically investigated the role of B cell expressed ion channels during KSHV lytic reactivation. Using known pharmacological ion channel modulators, genetic silencing approaches and electrophysiological analysis, we reveal for the first time that KSHV requires a B-cell expressed voltage-gated K^+^ channel, K_v_1.3, to enhance lytic replication. We show that the KSHV RTA protein upregulates K_v_1.3 expression via indirect Sp1-mediated transactivation. Enhanced K_v_1.3 expression and activity led to hyperpolarisation of the B-cell membrane potential, initiating Ca^2+^ influx through a T-type Ca^2+^ channel, Ca_v_3.2. T-type Ca^2+^ channels undergo both voltage-dependent activation and inactivation, with Ca_v_3.2 maximally inactivated at the depolarised membrane potentials that we measured from latently-infected cells^39^. The combination of reduced inactivation and the increase in driving force on Ca^2+^ ions resulting from hyperpolarisation, enhances Ca^2+^ influx via these channels. This led to Ca_v_3.2-mediated Ca^2+^ elevation and the subsequent Ca^2+^ driven nuclear localisation of NFAT1 to complete the KSHV lytic replication cycle. We therefore reveal the K_v_1.3-Ca_v_3.2 axis as a direct contributor to the lytic replication of KSHV in B cells (**Fig. 8**).

**Figure 8.**
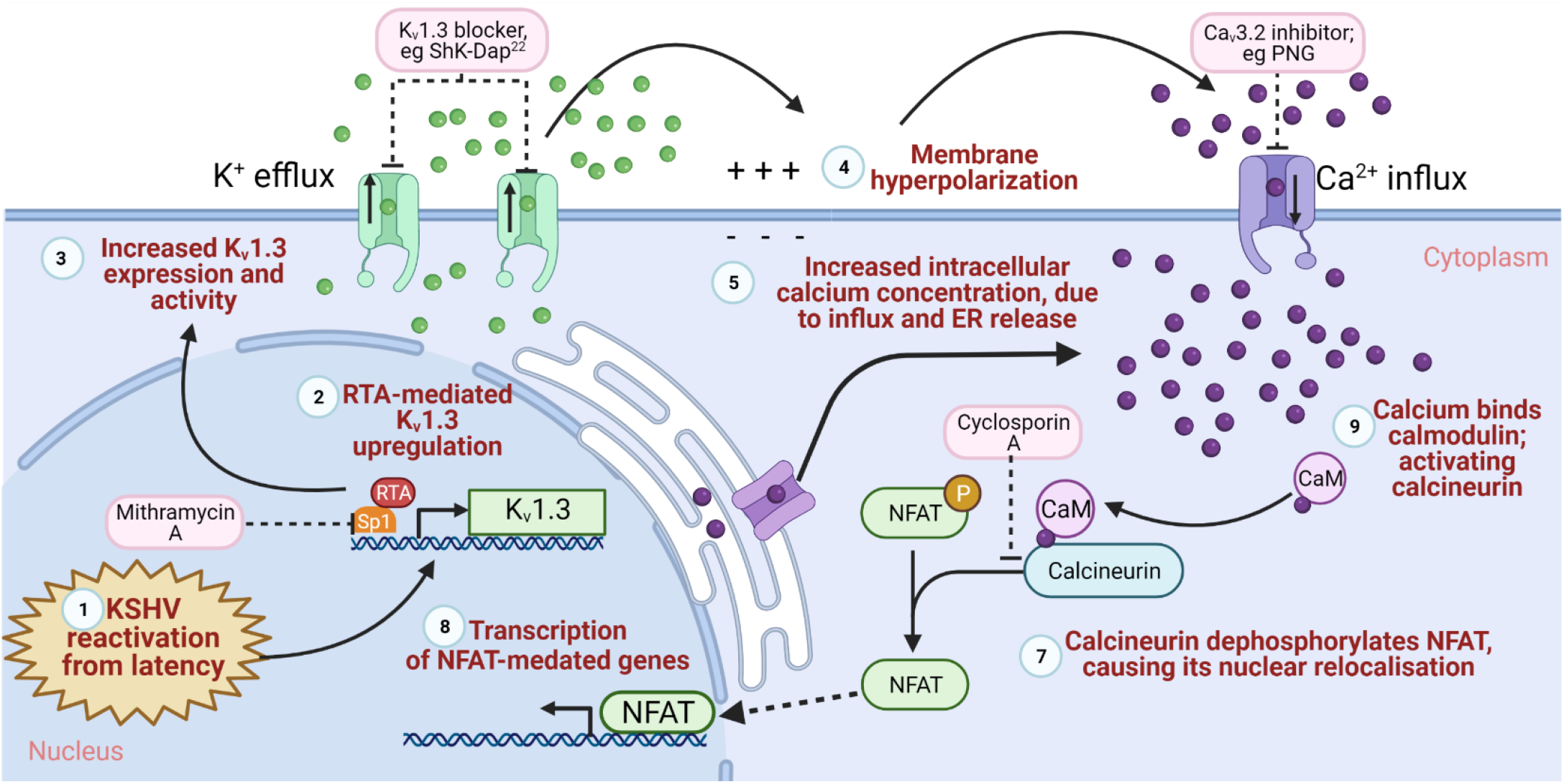
A K_v_1.3-Ca_v_3.2 axis is a direct contributor to KSHV lytic replication in B cells. Schematic representation of the KSHV-mediated K_v_1.3 induced hyperpolarisation and Ca_v_3.2-mediated calcium entry mechanism required for efficient lytic replication. Potential therapeutic interventions are highlighted. Created with Biorender.com

A striking feature of KSHV is the homology of its numerous ORFs to cellular genes^10^. These virus-encoded proteins contribute to KSHV-associated pathogenesis by subverting cell signalling pathways, including interferon-regulated anti-viral responses, cytokine-regulated cell growth, cell cycle progression and apoptosis. Many viruses encode viroporins^44^; ion channel proteins that modulate the ionic milieu of intracellular organelles to control virus protein stability and trafficking. However, no known viroporin exists amongst the ORFs of KSHV and it is therefore likely that evolution has tailor-made its proteins to regulate the expression of host cell ion channels to induce the Ca^2+^ signalling required for both latent and lytic replicative phases. Tumorigenesis may represent a by-product of this regulation, since in an array of human cancers, K_v_1.3 expression is enhanced and correlates with the grade of tumour malignancy^45^. It is also noteworthy that features of KS tumours mirror the phenotypic effects of K_v_1.3 overexpression, including the enhanced expression of inflammatory and angiogenic cytokines and uncontrolled cell cycle progression. This may reveal the KSHV driven activation of K_v_1.3 as a channelopathy, a group of diseases characterised by altered function of ion channel proteins or their regulatory subunits. Several ion channel inhibitors specifically target K_v_1.3, these inhibitors can either comprise small organic molecules such as quinine and 4AP or peptides purified from venom^37,46^. These venom-derived peptides are highly stable and resist denaturation due to the disulphide bridges formed within the molecules^46^. As with margatoxin, most are derived from scorpion venom, such as agitoxins, kaliotoxin, maurotoxin and noxiustoxin yet many inhibitors have been derived from ShK, a peptide originally isolated from the sea anemone *Stichodactyla helianthus^47^*. Given the abundance of natural sources for K_v_1.3-inhibition a safe, effective therapeutic based on these compounds is a promising target for prevention. Additionally, it is interesting to note that the anti-CD20 monoclonal antibody rituximab, a known K_v_1.3 inhibitor, substantially improves the outcome of KSHV patients^48^. Ca_v_3.2 inhibitors, including mibefradil, are also FDA approved as anti-hypertensive agents^49^, presenting additional opportunities for drug repurposing for the treatment of KSHV-associated diseases.

Finally, K_v_ channels have been previously identified as a restriction factor to the entry of both Hepatitis C virus^50^ and Merkel cell polyomavirus^51^, through their abilities to inhibit endosome acidification-mediated viral membrane fusion. Whilst the inhibition of endosomal acidification has been shown to reduce the entry and trafficking of KSHV virions, our electrophysiological analysis revealed enhanced cell surface K_v_1.3 activity during lytic replication that directly contributed to the hyperpolarised membrane potential of cells that was required for efficient KSHV replication. Thus, whilst additional roles of K_v_1.3 in endosomes cannot be excluded, our data suggest a divergent role of K_v_1.3 during herpesvirus infection that may be cell-type and/or virus specific.

In conclusion, we reveal the requirement of K_v_1.3 and Ca_v_3.2 for KSHV reactivation. To date, compounds targeting K_v_1.3 are in preclinical development^52^, whilst some Ca_v_3.2 blockers are FDA approved as antihypertensives and antiarrhythmics^49^. Further studies targeting these channels in KSHV-associated diseases are now warranted, which may reveal these channels as a host-directed therapeutic prospect for anti-KSHV drug development.

## Acknowledgements

This work was supported by a Rosetrees Trust PhD studentship, M662 (Whitehouse) and a Royal Society University Research Fellowship, G:480764 (Mankouri). We thank Professor Jae Jung, University of Southern California School of Medicine, Los Angeles, for the TREx BCBL1-RTA cells and Prof. J. Ladbury, University of Leeds for U87 cells and Dr Edwin Chen, University of Westminster for the lentivirus vectors. KSHV ORF59 and KSHV ORF65 antibodies were kind gifts from Prof. Britt Glaunsinger (University of California, Berkeley) and Prof. Shou-Jiang Gao (University of Pittsburgh), respectively.

