## Supplementary Figure 1 for "K_v_1.3 induced hyperpolarisation and Ca_v_3.2-mediated calcium entry are required for efficient Kaposi’s sarcoma-associated herpesvirus lytic replication"

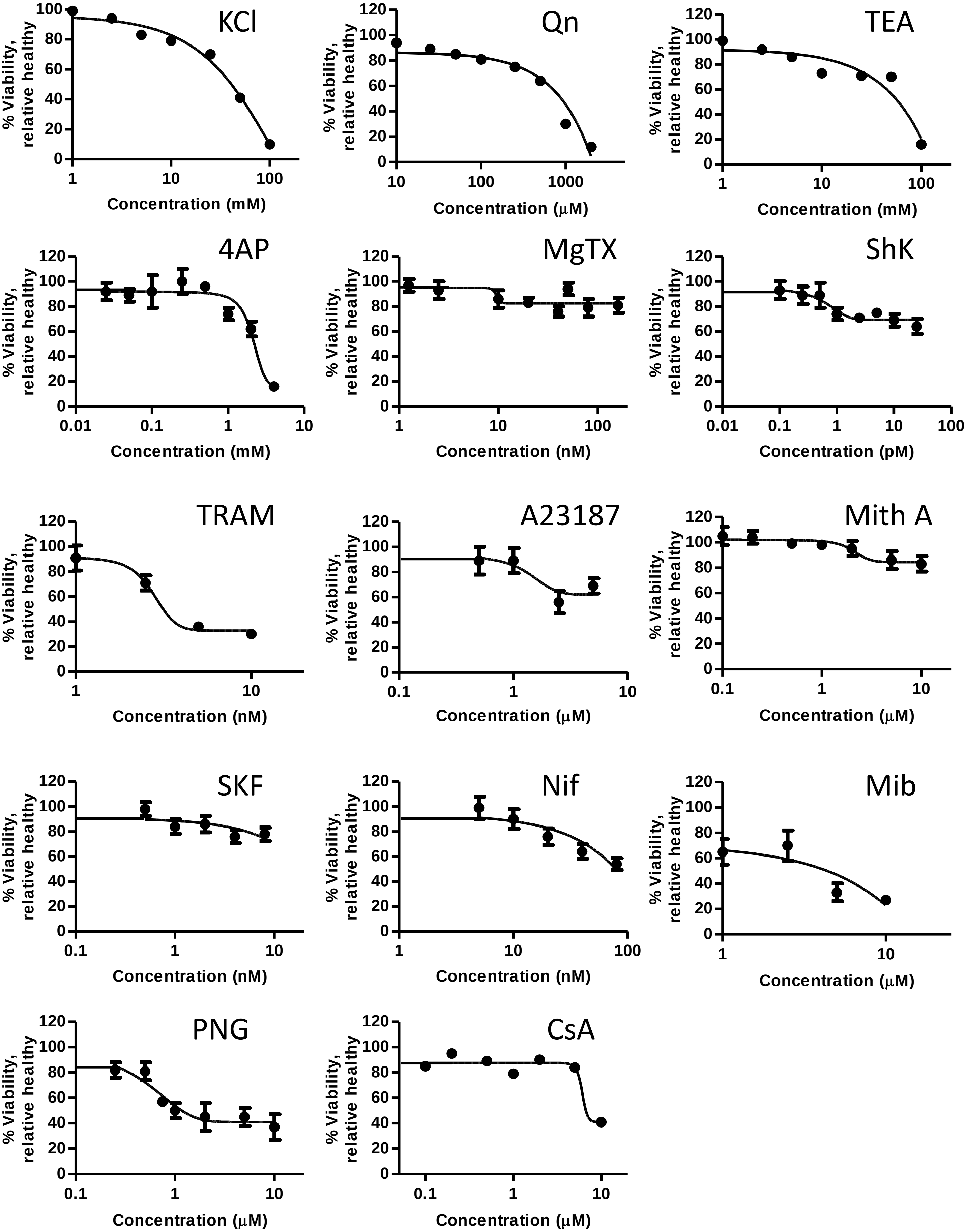


**Supplementary Figure 1.**

TREx BCBL1-RTA cells were seeded in triplicate in a flat 96-well tissue culture plates and treated with the indicated inhibitors for 24 h at a range of concentrations. Cell viability was measured using CellTiter 96 AQueous One Solution Reagent and absorbances measured at 490 nm.
